# Metagenomics and stable isotope probing offer insights into metabolism of polycyclic aromatic hydrocarbons degraders in chronically polluted seawater

**DOI:** 10.1101/777730

**Authors:** Ella T. Sieradzki, Michael Morando, Jed A. Fuhrman

**Affiliations:** Department of Biological Sciences, University of Southern California, CA, USA

## Abstract

Bacterial biodegradation is a significant contributor to remineralization of polycyclic aromatic hydrocarbons (PAHs): toxic and recalcitrant components of crude oil as well as byproducts of partial combustion chronically introduced into seawater via atmospheric deposition. The Deepwater Horizon oil spill demonstrated the speed at which a seed PAH-degrading community maintained by low chronic inputs can respond to an acute pollution. We investigated the diversity and functional potential of a similar seed community in the Port of Los Angeles, a chronically polluted site, using stable isotope probing with naphthalene, deep-sequenced metagenomes and carbon incorporation rate measurements at the port and in two sites further into the San Pedro Channel. We show a switch in the composition of the PAH degrading community from diverse early-responding generalists to late-blooming specialized degraders. This switch demonstrates the ability of the local seed community of degraders at the Port of LA to incorporate carbon from PAHs independently of a labile-hydrocarbon degrading succession. We were able to directly show that assembled genomes belonged to naphthalene degraders by matching their 16S-rRNA gene with experimental stable isotope probing data. Surprisingly, we did not find a full PAH degradation pathway in any of those genomes and even when combining genes from the entire microbial community. We use metabolic pathways identified in those genomes to generate metagenomic-based recommendations for future optimization of PAHs bioremediation.

## Introduction

Polycyclic aromatic hydrocarbons (PAHs) are recalcitrant, mutagenic and carcinogenic components of crude oil as well as byproducts of incomplete combustion [1]. Microbial biodegradation has an important role in PAH remediation alongside physical weathering processes [2]. Biodegradation of PAHs captured much scientific attention after the Deepwater Horizon (DWH) oil spill in the Gulf of Mexico in 2010. Several studies measured PAH degradation rates [3, 4] and showed enrichment of known PAH-degrading bacteria in beaches, surface water, the deep-sea plume and sediments even months after the spill began [5–9]. Bacteria known to have the ability to utilize PAHs as a carbon source include strains of *Cycloclasticus, Colwellia, Pseudomonas, Alteromonas*, and others [6, 10–12]. Many coastal sites worldwide experience chronic input of PAHs, mainly from atmospheric deposition and natural oil seeps. Recent studies show that chronic pollution supports a consistent “seed” of PAH degrading bacteria which can respond quickly to an acute pollution such as an oil spill [6, 10–15].

Stable isotope probing (SIP) is a well-established method for the identification of environmental bacteria utilizing targeted substrates, such as PAHs [16, 17], in which ^13^C PAHs are added to samples in order to make the DNA of PAH-utilizers heavy and thus capable of physical separation in a density gradient. A large-scale SIP study was performed on DWH surface and deep-plume water, revealing local strains of PAH-degrading bacteria that responded to the input of hydrocarbons [6]. Some genomes of those bacteria were assembled from mesocosm metagenomes in order to further explore their PAH metabolism [18]. The studies mentioned above, as many others, focus only on the high-density (i.e. most heavily ^13^C-labeled) fractions under the assumption that the most heavily labeled organisms, and thus the main targets, will be found there. However, this strategy may lead to overlooking degraders with low-GC (i.e. naturally lower DNA density) genomes, whose DNA may not appear in the heaviest fractions even if they include moderate amounts of ^13^C. Tag-SIP is a powerful and particularly sensitive extension of the standard SIP approach, in which DNA from both labeled samples and parallel unlabeled controls are separated into density fractions, and the 16S-rRNA coding gene is amplified from all density fractions of both samples for comparison. This approach, circumventing GC-based bias, allows us to track substrate incorporation by a single taxon demonstrated by an increase (shift) in its DNA density in the labeled samples compared to controls [19, 20].

One of the main motivations to study PAH-degrading organisms is to characterize their metabolic requirements. Understanding the suite of nutrients and cofactors those organisms require could potentially be applied towards bioremediation and biostimulation [21]. The combination of SIP with metagenomics can help reveal metabolic dependencies within assembled genomes of PAH-degraders [2, 15, 18, 21]. Additionally, it has been proposed that PAH biodegradation is a community process rather than fully performed by a single taxon [18]. Thus, it is important to investigate potential degradation using Tag-SIP and deep sequencing so as to identify not only the function of the organisms that initiate the degradation but also of the other organisms involved in later steps of degradation.

While the Port of LA (POLA) is not routinely monitored for PAHs, it is located in an area with natural oil seeps, it houses a ship-refueling station of high-aromatic-content marine diesel (ETC Canada, 11/22/2017) and it is surrounded by the second largest metropolitan area in the USA, which is likely a source of consistent atmospheric PAH deposition. The LA-Long Beach port is the busiest port in the United States and the 10^th^ busiest in the world according to the international association of ports and harbors (IAPH, 11/26/2017). A report published in 2010 revealed detectable levels of many PAHs at various stations within the port and as far as outside the port entrance, corresponding to our sampling site [22, 23]. Due to the high marine traffic at POLA, there is constant resuspension of the sediment into the water column, which would normally serve as a sink for PAHs due to their tendency to attach to particles [22, 23].

Here we provide evidence for the existence of a rare seed PAH-degrading, free-living (0.2-1 μ) microbial community in the chronically polluted Port of Los Angeles. We show that this seed can “spring into action” within 24 hours of high PAH input, and start a degradation process initially dominated by a variety of generalist taxa and later transformed into a specialized community dominated by two genera. We then discuss metabolic characteristics of the local PAH-degraders using largely complete genomes of PAH degraders and propose targets for biodegradation optimization experiments.

## Materials and Methods

### Sample collection

Surface seawater was collected on July 15^th^, 2014, October 6^th^, 2014 from three sites across the San Pedro Channel near Los Angeles, CA, USA (fig. 1): the port of Two Harbors, Santa Catalina Island (CAT), the San Pedro Ocean Time-series (SPOT) and the Port of Los Angeles (POLA). An additional sample was collected on May 15th, 2015 only from POLA. Water was collected by HDPE bucket into two 10 liter LDPE cubitainers and stored in a cooler in the dark until arrival to the lab.

**Figure 1:**
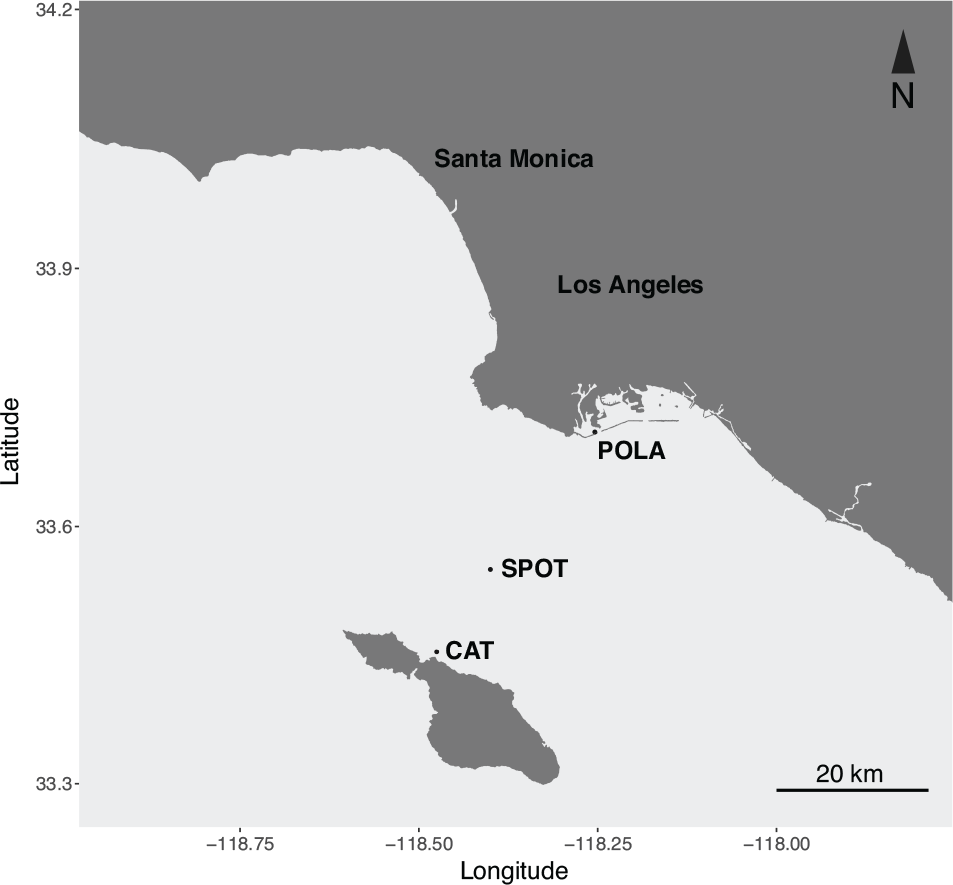
map of the sampling sites across the San Pedro Channel. The sites are within a range of 40 km

### Isotope addition and incubation

400 nM of either unlabeled (^12^C) naphthalene or fully labeled ^13^C-10-naphthalene (ISOTEC, Miamisburg, OH, USA) were added to separate 10 L bottles. This concentration is roughly 3 orders of magnitude lower than the solubility of naphthalene in water (30 mg/L = 234μM). In July and October, naphthalene-enriched seawater was incubated in the light in ambient temperature (17°C) for 24 hours, whereas in May it was incubated in the dark at ambient temperature (20°C) for 88 hours. The ambient temperature is similar to that measured in surface seawater (18.4°C in July and 19.9°C in May). At the end of each incubation period the seawater was filtered through an 80 nM mesh and a glass fiber Acrodisc (Millipore-Sigma, St. Louis, MO, USA) prefilter (pore size 1 μm) followed by a 0.2μm polyethersulfone (PES) Sterivex filter (Millipore-Sigma) to capture only the free-living microbes. After filtration, 1.5ml Sodium-Chloride-Tris-EDTA (STE; 10mM Tris-HCl, 1mM EDTA and 100mM NaCl) buffer was injected into the Sterivex casing and the filters were promptly sealed and stored in −80°C.

### Carbon incorporation rate measurement

In October 2014, seawater from all three sites was incubated in quadruplicate 2-liter polycarbonate bottles with 400nM ^13^C labeled naphthalene. 1μM ammonium-chloride was also added to each bottle to prevent nitrogen limitation. A single bottle was filtered immediately after amendment addition to establish a t_0_ atom% ^13^C of the particulate carbon at each site. The remaining water was incubated in a temperature-controlled room (see above). Incubations were carried out for ~24 h and were terminated by filtration onto pre-combusted (~5 h at 400°C) 47mm GF/F filters (Whatman, Maidstone, United Kingdom). The filters were then dried at 60°C and kept in the dark until analysis. Isotopic enrichment was measured on an IsoPrime continuous flow isotope ratio mass spectrometer (CF-IRMS). IRMS data were corrected for both size effect and drift before being calculated as previously described (Dugdale and Goering, 1967).

### DNA extraction

Total DNA was extracted from the Sterivex filters using a modified DNeasy Plant kit protocol (Qiagen, Hilden, Germany). The Sterivex filters containing STE buffer were thawed. 100 μL 0.1mm glass beads were added into the filter casing and put through two 10-minutes cycles of bead beating. The flow-through, pushed out using a syringe, was incubated for 30 minutes at 37°C with 2 mg/ml lysozyme followed by another 30 minutes incubation at 55°C with 1 mg/ml proteinase K and 1% SDS. The resulting lysate was loaded onto the DNeasy columns followed by the protocol as described in the kit instructions. Only samples from POLA 5/2015, POLA 7/2014 and SPOT 10/2014 yielded enough DNA for ultracentrifugation of both the labeled and unlabeled DNA. However, metagenomes were sequenced from all samples other than SPOT 10/2014.

### Ultracentrifugation and density-fractions retrieval

Isopycnic ultracentrfugation and gradient fractionation were performed as described in previous work [20, 24]. Briefly, DNA from the labeled and unlabeled samples was added into separate quick seal 5ml tubes (Beckman Coulter, Indianapolis, IN, USA) combined with 1.88 g/ml CsCl and gradient buffer for a final buoyant density of 1.725 g/ml. The tubes were sealed and centrifuged in a Beckman Optima L100 XP ultracentrifuge and near-vertical rotor NVT 65.2 at 44100 rpm at 20°C for 64 hours.

The gradient was divided into 50 fractions of 100μl each. Refraction was measured using 10 μl of each fraction using a Reichert AR200 digital refractometer, and converted into buoyant density (ρ=10.927*n_c_−13.593) [25]. DNA in each fraction was preserved with 200 μl PEG and 1 μl glycogen, precipitated with ethanol, eluted in 50 μl Tris-EDTA (TE) buffer and quantified using PicoGreen (Invitrogen, Carlsbad, CA, USA).

### Amplification of the 16S-rRNA V4-V5 hypervariable regions

PCR was performed on each fraction with detectable DNA. Each reaction tube contained 12 μl 5Prime Hot Master Mix (Quantabio, Beverly, MA, USA), 1 μl barcoded 515F-Y forward primer (10 μM), 1 μl indexed 926R reverse primer (10 μM), 1 ng of DNA and 10μl molecular-grade water.

Thermocycling conditions:

1. 3 min denaturation at 95°C
2. 30 cycles of denaturation at 95°C for 45 sec, annealing at 50°C for 45 sec, and elongation at 68°C for 90 sec
3. final elongation at 68°C for 5 min

PCR products from each fraction were cleaned using 1x Agencourt AMPure XP beads (Beckman Coulter), quantified by PicoGreen and diluted to 1 ng/μl. A pool of 1 ng of each uniquely-barcoded product was cleaned and concentrated again with 0.8x Agencourt AMPure XP beads. The pooled amplicons were sequenced on Illumina MiSeq (UC Davis, USA) for 600 cycles. Each pool was also spiked with 1ng of an even and a staggered mock community in order to assess the sequencing run quality [26]. All expected OTUs were found in the observed mock communities and accounted in total for 99.5% of the reads, indicating that there was no unexpected bias in the sequencing run [27].

### Amplicon data processing

The raw reads were quality-trimmed using Trimmomatic [28] version 0.33 with parameters set to LEADING:20 TRAILING:20 SLIDINGWINDOW:15:25 MINLEN:200 and merged with Usearch version 7 [29] with a limit of maximum 3 differences in the overlapping region. The resulting merged-reads were analyzed in Mothur [30] and clustered at 99% identity following the MiSeq SOP (March 20^th^, 2016) [31] with one exception: we found that degapping the aligned sequences, aligning them again and dereplicating again fixed an artifact in which a few abundant OTUs were split due to the alignment, despite 100% identity, due to an additional terminal base between the OTU representative sequences. OTUs with a total of less than 10 reads over all fractions were removed from the analysis. The remaining 2366 OTUs were assigned taxonomy using the arb-silva SINA search and classify tool version 1.3.2 [32].

### Detection of OTU enrichment due to substrate incorporation

Plots of normalized abundance as a function of density were generated in R (https://www.r-project.org/) for the top 200 most abundant OTUs in each sample. The abundance of each OTU was normalized to a sum of 1 across all fractions (eq. 1).

Equation 1: Normalization of OTU abundance per fraction where i=fraction number, Ri=relative abundance of the OTU in fraction i, Di=DNA quantity in fraction i, Ni=normalized abundance of the OTU per fraction i

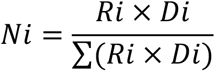

To detect enrichment, the weighted mean density of an OTU in the labeled and unlabeled samples was calculated and if the difference exceeded 0.006 g/ml the shift was determined real. This is a conservative threshold chosen to reduce the potential for false positives. Only abundant, non-spurious OTUs led to plots that were visually validated as consistent enough to calculate enrichment. With these criteria, only 88 validated OTUs from July 2014 and 127 from May 2015 were further analyzed.

### Metagenomic library preparation

The original unfractionated DNA extracted from the Sterivex filters was sheared by Covaris m2 to a mean length of 800 bp. Libraries were prepared from 15 ng of sheared DNA using the Ovation Ultra-low DR Multiplex System v2 kit (Nugen, Redwood City, CA, USA) with 9 amplification cycles. The libraries were bead purified as described above and sequenced on Illumina MiSeq for 600 cycles (UC Davis, USA) or on Illumina HiSeq rapid run for 500 cycles (USC genome core). See sup. table S1 for a detailed list of sequenced metagenomes.

### Metagenomic sequencing analysis

Reads were quality-trimmed using Trimmomatic version 0.33 as described above. Paired reads were assembled per sample in an iterative subsampling and assembly process as described in Hug et al. [33] but using metaSPAdes version 3.9.1 instead of IDBA-UD, followed by overlap assembly with minimus 2 with minimum overlap 200 bp and minimum identity 99%. Paired reads from all sequenced samples (sup. table S1) were mapped back to the contigs with BBmap (sourceforge.net/projects/bbmap/) requiring 95% identity. Binning was performed using two approaches: (1) binning the POLA 5/15 13C metagenome with CONCOCT [34] and bin refinement in Anvi’o [35]. (2) pooling contigs longer than 5kbp from all four naphthalene-enriched metagenomes, dereplicating them with cd-hit at 99% id [36, 37] and binning them using a combination of MaxBin2 [38], CONCOCT [34] and MetaBAT2 [39]. These bins were combined, refined and reassembled using the MetaWRAP pipeline [40]. Genomic bins generated by both methods were dereplicated using dRep [41] and only bins that were at least 50% complete and under 10% redundant were analyzed.

Initial taxonomic assignment of MAGs was performed using GTDB-Tk [42]. We then improved the taxonomy by generating class-level phylogenomic trees with GToTree [43] using NCBI RefSeq complete genomes and placing the bins assigned to the class by GTDB-Tk within them.

### Metabolic analysis

Open reading frames (ORFs) in the final set of metagenomic assembled genomes (MAGs) and viral contigs were predicted using Prodigal [44] and annotated by assignment to clusters of orthologous groups (COGs) using the Anvi’o anvi-run-ncbi-cogs function. KEGG (Kyoto encyclopedia of genes and genomes) orthology for ORFs was assigned with Kofamscan using the prokaryote.hal profile and its built-in thresholds [45]. The results were then visualized using the R package Pathview [46]. Additionally, ORFs were searched against NCBI NR database using DIAMOND [47] and the best hits as well as KEGG annotations are detailed in sup. table S2. Kofamscan results were also summarized using KEGGdecoder [48] (sup. fig. S1). ORF taxonomy was determined by Kaiju [49] using the RefSeq database.

Read recruitment from different samples to the MAGs and viral contigs was visualized with Anvi’o [35] using the Q2Q3 view. This setting ignores the 25% lowest covered and 25% highest covered positions within the MAG when calculating mean coverage to avoid bias due to islands or highly conserved genes.

### Viral contigs

523 viral contigs were identified using VirSorter version 1.0.3 [50] with the Virome database. Another 20 viral contigs were added by running VirFinder version 1.1 [51] with the Tara Oceans training set, predicting ORFs with prodigal, searching the protein sequences against GenBank’s non-redundant database and keeping only contigs with at least two best hits to viruses.

### Data availability

Metagenomic and amplicon raw reads from enrichment mesocosms can be found on EMBL-ENA under project project PRJEB26952, samples ERS2512855-ERS2512864. The metagenomic library blank is under sample ERS2507713. Amplicon reads can be found under sample accession numbers ERS2507470-ERS2507679, PCR blanks and mock communities under samples ERS2507702-ERS2507712. Metagenomic t0 raw reads can be found under project PRJEB12234, samples ERS2512914-ERS2512919.

## Results

### Naphthalene degradation rates across the San Pedro Channel

Our main hypothesis was that at the Port of Los Angeles (POLA) there would be a seed of PAH-degrading bacteria due to constant, chronic inputs. We also wanted to test whether this degradation potential extended out into the San Pedro Channel at the San Pedro Ocean Time-series (SPOT) and Two Harbors (CAT) (fig. 1). Indeed, isotopic enrichment measurements at POLA indicated a mean naphthalene uptake rate of 35 nM/d (standard deviation 19.53 nM/d) given a high input of 400nM naphthalene. However, naphthalene incorporation rates at SPOT and CAT were below detection.

### SIP-identified PAH-degrading taxa at POLA

Naphthalene-enrichment of POLA seawater was performed twice: July 2014 and in May 2015. The incubation experiments differed in duration (24 hours in July and 88 hours in May) and in light availability (10% ambient light in July and dark in May). In both cases we observed taxa that incorporated labeled carbon. However, the diversity of operational taxonomic units (OTUs) with significant enrichment (>0.006 g/ml buoyant density increase corresponding to 11 atom %excess) in May was lower than in July (table 1, fig. 2) and the main degraders in May, *Colwellia* sp. and *Cycloclasticus* sp. were not detected in July. Significant density shifts were detected in 56 out of 84 OTUs in July and in 6 out of 120 OTUs in May. Of the naphthalene incorporator OTUs, 26 were highly enriched (>0.009 g/ml buoyant density increase corresponding to 16.5 atom %excess) in July and 3 were highly enriched in May.

**Table 1:** Taxonomic distribution of the number of total and significant incorporators OTUs at POLA in July 2014 and May 2015. In July (24h incubation) both richness and diversity of significantly enriched OTUs were high, whereas in May almost all enriched OTUs were Gammaproteobacteria.

| Taxonomic group |
| --- |
| Alphaproteobacteria |
| Betaproteobacteria |
| Gammaproteobacteria |
| Unclassified Proteobacteria |
| Flavobacteriia |
| **Opitutae** |
| **Verrucomicrobia** |
| **Cytophagia** |
| **Sphingobacteriia** |
| **Deferribacteres** |
| **Acidimicrobiia** |
| **Planctomycetes** |
| **Photosynthetic eukaryote** |
| **total** |

**Figure 2:**
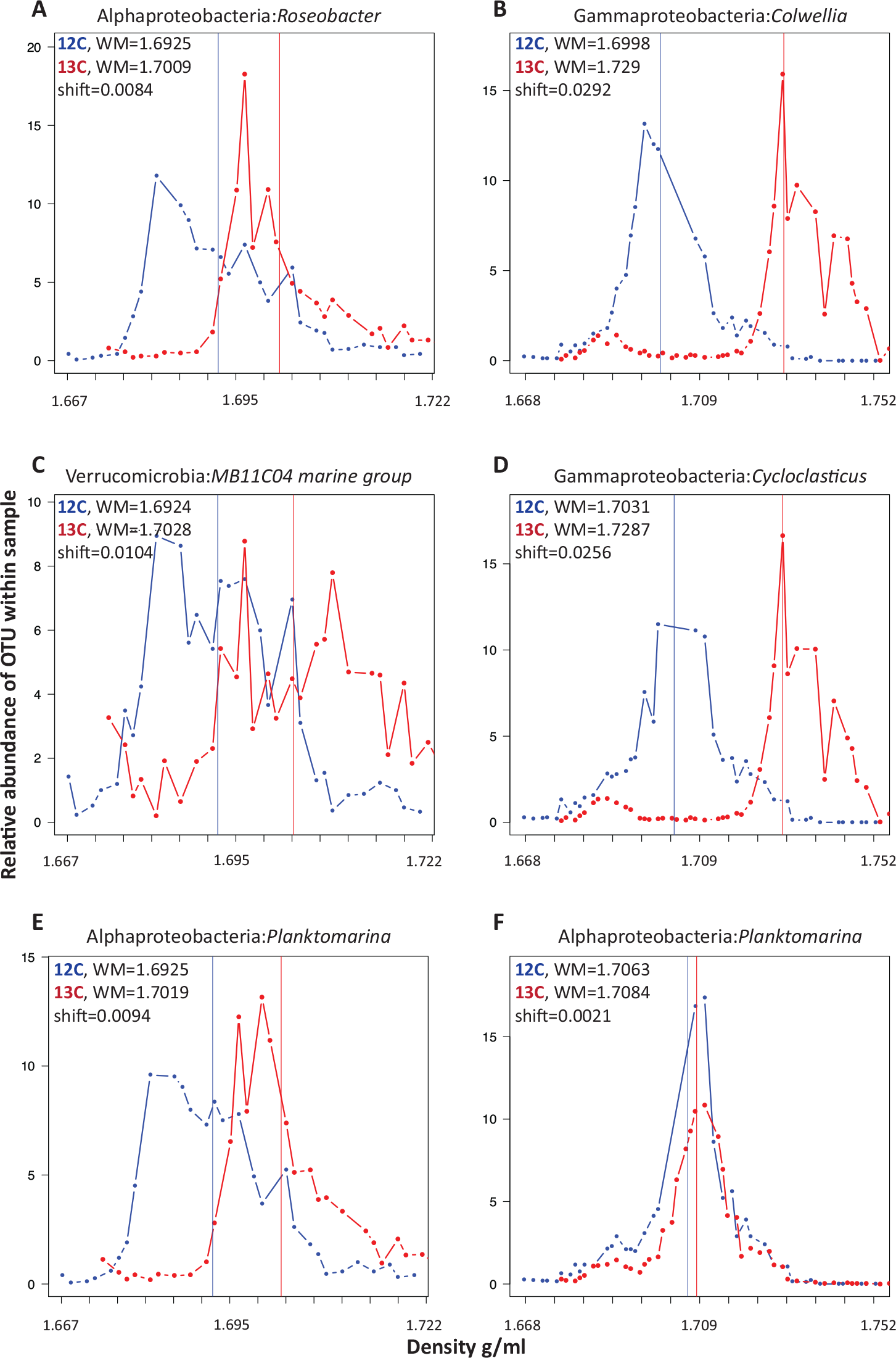
Distribution of labeled (13C, red) and control (12C, blue) relative abundance of specific OTUs as a function of density from POLA samples in July 2014 (left; A,C,E) and May 2015 (right; B,D,F). Vertical lines represent the weighted mean of the distribution. The weighted mean density (WM) of the labeled (^13^C) and control (^12^C) and the difference between (density shift) them are noted on each plot.

The main naphthalene degraders identified in May were rare (<0.1% relative abundance) to non-detectable prior to enrichment (t_0_) at all sites in all dates. However, after 88 hours of incubation *Colwellia* sp. and *Cycloclasticus* sp. (Gammaproteobacteria) comprised 40% of the free-living (0.2-1 μ) microbial community.

Naphthalene enrichment of SPOT seawater in October 2014 yielded only three OTUs with significant shifts belonging to the clades: Marine Group A (SAR406), Rhodospirillaceae (Aegean-169) and Flavobacteriia NS5 marine group. Enrichment of all these OTUs was marginal (0.0062, 0.0061 and 0.0066 g/ml buoyant density increase respectively). However, a trend appeared in many OTUs with enrichment lower than 0.006 g/ml in which there was a bimodal distribution of the OTU normalized abundance where one distribution corresponded with the unlabeled OTU and the other was centered around a higher density (sup. fig. S2), implying that only a part of the population incorporated naphthalene-originated carbon.

Relative abundance of the most abundant subphyla between t_0_ and 24 hours of incubation with naphthalene were similar between POLA and SPOT. In both sites there was an increase in Gammaproteobacteria and a decrease in Flavobacteriia and chloroplasts representing photosynthetic eukaryotes (sup. fig. S3). At POLA after 88 hours the change was even more extreme, with Gammaproteobacteria abundance increasing further at the expense of the normally dominant Alphaproteobacteria.

Breakdown of these patterns at a higher taxonomic resolution revealed that the observed increase in Gammaproteobacteria was mainly due to proliferation of *Colwellia* and *Cycloclasticus* OTUs. These OTUs were enriched at 53 (Colwellia) and 47 (Cycloclasticus) atom %excess [52] (fig. 2B,D). While both taxa were represented by multiple OTUs, there was always a dominant OTU which matched the 16S-rRNA genes from the MAGs (see below). The most abundant OTU accounted for 95% and 99% of *Colwellia* and *Cycloclasticus* amplicons, respectively.

### Metagenome-assembled genomes (MAGs)

We assembled and binned 43 dereplicated MAGs that were more than 50% complete (mean=88%, SD=11%) and less than 10% redundant from naphthalene-enriched POLA water (mean=4.6%, SD=2.6%). We then mapped reads from naphthalene-enriched and unenriched (t_0_) metagenomes to those bins in order to pinpoint potential degraders, under the assumption that potential degraders would be more abundant in mesocosm metagenomes compared to t_0_ metagenomes. Two MAGs of interest were classified as *Colwellia* sp. and *Cycloclasticus* sp. and contained 16S-rRNA genes that matched at 100% identity to the most abundant OTUs of PAH-carbon incorporators.

The *Colwellia* MAG (3.54 Mbp; 91% complete; 3.6% redundant) had high coverage and breadth (portion of the bin that has at least 1x coverage) only in naphthalene-enriched POLA water from May and October (fig. 3 A). Its closest relative based on a phylogenomic tree of 117 single-copy genes [43] is *Colwellia* sp. PAMC 20917 isolated from the Mid-Atlantic Ridge cold, oxic subseafloor aquifer [53] (sup. fig. S4; sup. table S3). This MAG contained subunit B of naphthalene dioxygenase gene, which is the first step in naphthalene degradation, but only parts of the remainder of the pathway known from pure cultures (sup. table S2). In addition. the MAG contained near-complete chemotaxis and flagella-assembly, near complete vitamin B6 and biotin biosynthesis, and a complete riboflavin biosynthesis pathway. This organism has transporters for nitrite/nitrate, urea, phosphate, molybdate and heme. Finally, it can transport nitrate and potentially reduce it to ammonium via a dissimilatory pathway (sup. table S2; S4).

**Figure 3:**
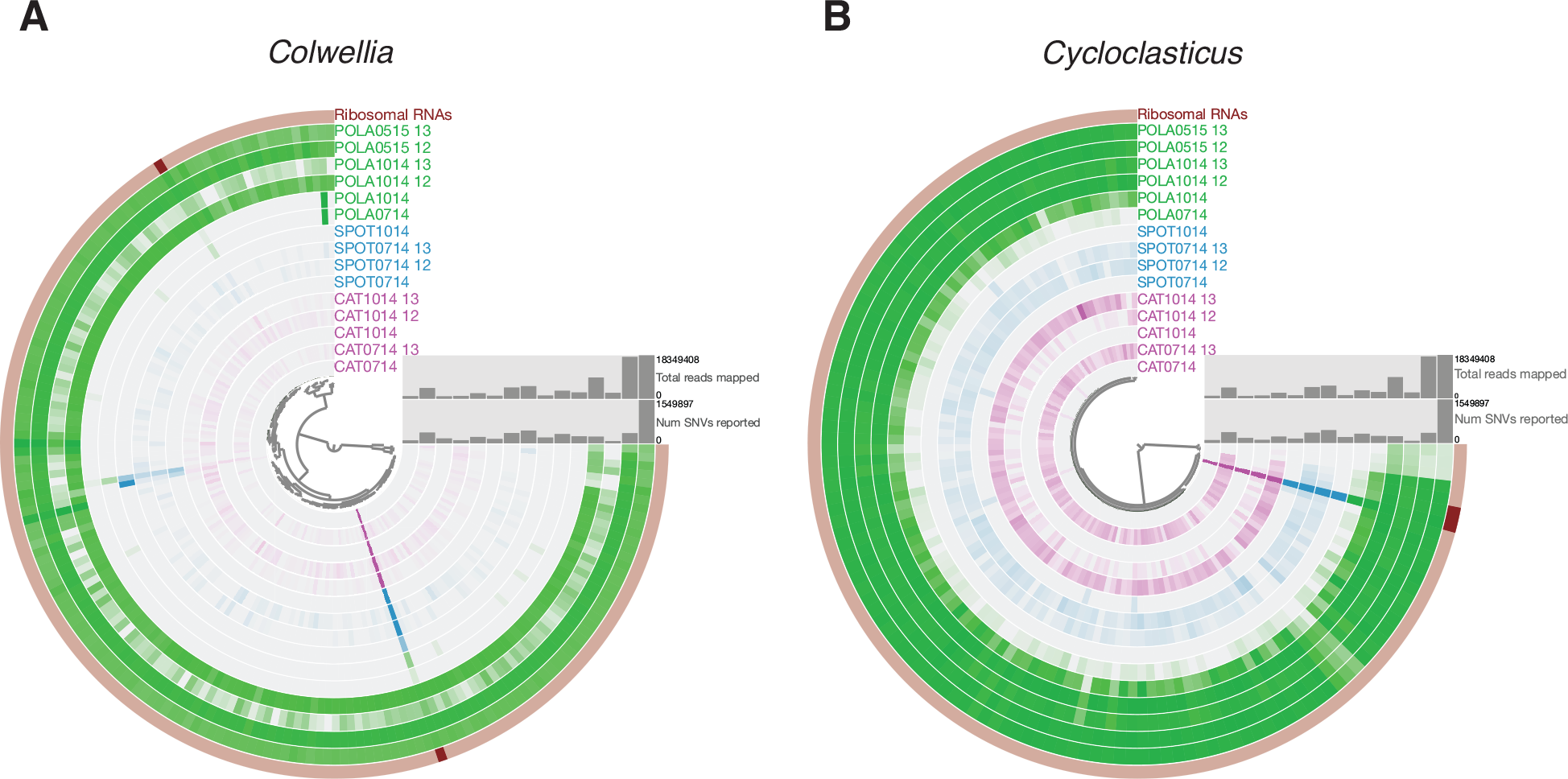
Mean coverage of (A) *Colwellia* and (B) *Cycloclasticus* MAGs across sites, dates and naphthalene enrichment. Each concentric circle represents one sample. Each bar across the concentric circles denotes one contig within the MAG. The bar color intensity corresponds to mean coverage of the contig within a sample. The central tree shows hierarchical clustering of the contigs by tetranucleotide frequency and mean coverage. Enriched mesocosms have 12 (^12^C-naphthalene added) or 13 (^13^C-naphthalene added) next to the sample name. The other samples represent t0.

The *Cycloclasticus* MAG (99% complete; 2.9% redundant, 2.4 Mbp) was detected in POLA naphthalene-enriched metagenomes from May, July and October (fig. 3 B). This MAG was most closely related to *Cycloclasticus zancles* 78-ME (sup. fig. S4; sup. table S3). This organism has a near-complete flagellar assembly pathway but not the chemotaxis pathway. It can incorporate nitrogen from cyanate and biosynthesize riboflavin and biotin. It also includes complete degradation pathways for cymene (xylene) and benzene. Similar to the *Colwellia* MAG, this MAG also contains transporters for nitrite/nitrate, urea and phosphate, and can potentially reduce nitrate to ammonium (sup. table S2; S4).

In addition to *Colwellia* and *Cycloclasticus*, six more MAGs contained 16S-rRNA genes identified by a hidden Markov model (HMM) which matches enriched OTUs at >99% identity. We assigned taxonomy to these 16S-rRNA sequences using the arb-Silva SINA aligner [32] and compiled the best hits to OTUs from these MAGs in table 2. None of these MAGs contained the PAH or BTEX (benzene, toluene, ethylbenzene, xylene) degradation pathways defined by pure culture studies (sup. table S4). However, the *Porticoccus*, SAR92 and Rhodobacteraceae MAGs contained a biphenyl dioxygenase gene, the *Puniceispirillum*, Rhodobacteraceae and *Pseudohongiella* MAGs included an aromatic ring-hydroxylating dioxygenase (sup. table S2). Three of these MAGs can assimilate nitrogen from nitroalkanes (fuel additives) or nitriles. Additionally, we assembled 26 medium (50-70%) to high (>70%) completion MAGs which did not contain 16S-rRNA genes but were taxonomically classified to groups with enriched OTUs (table 3), indicating they may have PAH-degradation pathways or pathways involved in downstream mineralization (e.g. catechol degradation or single aromatic ring dioxygenase). General metabolic pathways of all MAGs are summarized in a heat map (sup. fig. S1). Since several studies suggested that full degradation of PAHs may be performed by bacterial consortia [18, 54], we also investigated the metabolic potential of the whole assembled community (contigs >= 1kbp). The full naphthalene degradation pathway identified in pure cultures and incorporated into the KEGG database was not found.

**Table 2:** MAGs of potential naphthalene degraders identified by assembled 16S-rRNA identity to enriched OTUs. High-completion MAGs are highlighted. See sup. table S5 for MAG coverage per sample.

| OTU taxonomy/ID | MAG ID | 16S-rRNA identity between MAG and OTU | Alignment length (bp) | MAG Completion (%); redundancy (%); size (Mbp) |
| --- | --- | --- | --- | --- |
| *Cycloclasticus*/3 | 42_1 | 100% | 372 | 99.3; 2.9; 2.4 |
| *Colwellia*/1 | POLA0515-13_bin_34 | 100% | 372 | 91.4; 3.6; 3.5 |
| *Litoricola*/41 | 27_2 | 100% | 373 | 100; 7.2; 2.2 |
| *Pseudohongiella*/36 | POLA0515-13_bin_4 | 100% | 373 | 61.2; 4.3; 3.8 |
| Rhodobacteraceae/118 | POLA0714-12_bin_12 | 100% | 372 | 97.8; 1.4; 2.9 |
| *Puniceispirillum*/13 | 3_1 | 99.5% | 371 | 90.7; 2.9; 1.6 |
| SAR92/16 | POLA0515-13_bin_1 | 99.5% | 373 | 68.4; 0.7; 2.3 |
| *Porticoccus*/25 | 47_1 | 100% | 373 | 97.1; 0.7; 1.2 |

**Table 3:** MAGs of potential PAH degraders determined by general MAG taxonomy and enriched OTU taxonomy. The rightmost column is the number of OTUs in the taxonomic group of the MAG which were not matched to a MAG containing 16S-rRNA.

| Taxonomic group | Number of MAGs without 16S-rRNA | Number of enriched OTUs without a matching MAG |
| --- | --- | --- |
| Rhodobacteraceae | 7 | 7 |
| Gammaproteobacteria | 11 | 17 |
| Alphaproteobacteria | 8 | 16 |

Moreover, to confirm we did not miss additional naphthalene degradation genes because they did not assemble into contigs, we searched all forward reads from the POLA May 2015 ^13^C metagenome against all nucleotide sequences of naphthalene dehydrogenase ferredoxin subunit (nahAc, the first step of naphthalene degradation) from NCBI GenBank (188 references). We found 248 reads (0.0008% of the total metagenome) that hit with a relaxed cutoff of e-value 10^−3^ (mean identity 97.8%, minimum bitscore 30). Using a conservative estimate that this gene comprises 0.1% of the metagenome (similar to 16S-rRNA), assuming one copy per cell and that each read maps to one copy of the gene, this would imply that only 8 out of 10,000 cells carry the naphthalene dioxygenase nahAc gene. This fraction would decrease even further if we dropped the single-copy and one read per gene assumptions. With that, many reads map to other dioxygenase enzymes related to degradation of aromatic compounds (e.g. homogentisate 1,2 dioxygenase, catechol 1,2 dioxygenase).

### Viral community

The assembled contigs from POLA naphthalene-enriched microcosms included contigs that were recognized as viral using two tools: (1) VirSorter [50] which relies on identification of hallmark viral genes as well as depletion of Pfam proteins and lack of strand switching, and (2) VirFinder [51] which relies on k-mer frequencies and was trained on a curated set of marine bacteria and viruses from the TARA Oceans data [55]. Ignoring the less reliable VirSorter results (categories 3 and 6) we identified 586 viral contigs, 3 of them circular (15.7, 33.1 and 35.7 kbp), and 9 potential prophage contigs. In line with the bacterial community, these contigs represented a viral community unique to POLA, most of which was not detected in unenriched samples (sup. fig. S5).

## Discussion

### Naphthalene biodegradation potential does not extend outside the Port of Los Angeles

Naphthalene degradation rates were detectable only at POLA but not at SPOT or CAT. As there is tidal mixing across the San Pedro Channel, we would expect some potential for degradation at those sites by bacteria advected from the port. A possible explanation for the non-detectable rates at SPOT and CAT is that diminishing PAH inputs offshore [56] can support a very limited degrader seed community, and that the PAH degraders are out-competed for nutrients at those sites by microbes that are better adapted to oligotrophic conditions, and lose viability or are otherwise removed by processes like grazing and viruses at rates faster than they are replenished by mixing or growth. Tag-SIP results at SPOT indicated that only three OTUs were significantly enriched and others presented a bimodal distribution which may imply PAH incorporation only by a sub-population of the OTU. However, in those operational taxonomic units (OTUs) we did not observe enrichment of the same magnitude as that observed at POLA, which may support degradation rates too slow and concentrated in a small fraction of the community to be detected. More studies quantifying these rates across San Pedro Channel are needed to determine if this pattern continues throughout the year.. The PAH carbon incorporation rate at POLA, however, indicated a minimum removal of ~10% of the initial concentration per day. This rate is likely underestimated as measurement was performed using GF/F filters, which have a pore size leading to loss of roughly half of marine free-living prokaryotes. In addition, there could well be degradation of PAH without incorporation of the labeled carbon into cells.

### Naphthalene-degrading taxa vary by incubation conditions

Amplifying 16S-rRNA from all density fractions allows us to track specific strains, or operational taxonomic units (OTUs), that were enriched due to incorporation of carbon from naphthalene. Previous papers using PAH-SIP only analyzed the taxonomic composition of the highly-enriched density fractions [6, 18]. While this process still reveals major contributors to PAH degradation, it overlooks taxa with lower enrichment, which may use intermediate degradation products or additional carbon sources, but still participate in the PAH mineralization pathway. These taxa play a key role in the full degradation and detoxification of PAHs.

Incubation of POLA water with naphthalene revealed a difference in PAH degrading communities between July 2014 and May 2015. Several factors could contribute to the difference between those communities. Incubation in the dark in May promoted heterotrophy and may have minimized competition for nutrients with photoautotrophic organisms. However, if that was the case, we would expect a higher relative abundance of cyanobacteria and photosynthetic picoeukaryotes in July compared to May, which we did not observe. Secondly, the duration of the experiment may have contributed to the accumulation of labeled carbon in bacterial DNA. In order to observe pronounced enrichment, the PAH degraders have to replicate at least once, and DNA density increases with every replication as the new strand is made with labeled nucleotides. Additionally, it is methodologically difficult to observe enrichment in rare taxa [57]. Since the seed degrader community is made up of rare organisms, their abundance must increase substantially before their enrichment can be tracked. The other side of the coin is that a longer incubation increases the chance of cross-feeding, namely labeling of organisms that incorporate intermediate- or end-products of naphthalene degradation. The observed shifting of chloroplast OTUs is a prime potential example, though not easily explained. Incorporation of naphthalene-originated carbon into chloroplasts can imply several things: (1) that those chloroplasts belonged to mixotrophic algae that can also degrade recalcitrant hydrocarbons (2) that they belonged to mixotrophic algae that feed on PAH-degrading bacteria, or (3) that they are photoautotrophs that incorporated respired remineralized labeled carbon dioxide either through carbon fixation or through anaplerotic pathways. While few algae capable of partial naphthalene degradation or detoxification have been identified [58] (e.g. *Chlorella vulgaris*, *Dunaliella* sp., *Scenedesmus obliquus*, *Nitzschia* sp. and *Skeletonema costatum*), there is no evidence of incorporation of the resulting carbon into biomass [59–61]. Furthermore, the taxonomy assigned to the ^13^C-enriched chloroplasts was of picoeukaryotic genera *Micromonas*, *Bathycoccus* and *Ostreococcus*, none of which are known to have PAH-degradation capabilities, though the full characteristics of “wild” members of these genera are not understood. Phagotrophy of PAH-degrading bacteria also seems an unlikely explanation as picoeukaryotic mixotrophs have been shown to ingest only a few cells per hour [62]. Even if every cell they ingest was fully labeled, between this grazing rate, the rate of prey-originated carbon incorporation into nucleotides, the size of eukaryotic genomes and the time limitation of 24 hours for replication, significant enrichment seems unlikely. Though significant labeling of chloroplasts has been observed previously in 24 hour incubations within the San Pedro Channel [63], enrichment of chloroplasts due to cross-feeding of respired labeled carbon seems unlikely due to the dilution of labeled respired bicarbonate in the 2mM pool of unlabeled bicarbonate present in seawater [64], unless these eukaryotes are located in enriched microzones that are also PAH degradation hotspots (e.g. perhaps suspended particles).

Chloroplasts aside, cross-feeding of intermediate products of naphthalene degradation is a vital part of the complete remineralization of this hydrocarbon which appears to have been often overlooked.

A high input of PAHs can trigger a succession process that would affect the microbial community composition. It is likely that over time there is a shift from diverse generalist degraders that respond quickly and would be prominent after 24 hours to an extremely uneven community dominated by PAH-degrading specialists after several days. Indeed, we saw taxonomic diversity of OTUs whose density shifted after 24 hours, representing 8 phyla, as opposed to a community dominated by Gammaproteobacteria, mainly *Colwellia* and *Cycloclasticus*, in both relative abundance and richness of shifted OTUs. Guttierez et al. [6] also identified 4-5 orders of magnitude increase in abundance of *Colwellia* and *Cycloclasticus* in mesocosms after three days. Another driving force behind a quick succession may be high abundance and diversity of viruses [65, 66], as suggested by the abundance of assembled viral contigs and the strikingly unique viral community in enriched mesocosms in May. This viral community is potentially comprised of viruses that infected organisms that were at least moderately abundant during the 88 hours of incubation [67]

### Metabolism of putative PAH-degraders

In order to gain more insight into metabolic requirements of naphthalene degrading bacteria at POLA, we examined Kegg pathways and key transporters and enzymes within the annotated proteins in our metagenomic assembled genomes (MAGs). To date, only one published study characterized metabolic pathways in assembled genomes of marine hydrocarbon degrading bacteria, using metagenomes from naphthalene- and phenanthrene-enriched seawater from the Deepwater Horizon (DWH) oil spill [6, 18]. Within the DWH mesocosms, the prominent naphthalene degraders belonged to the genera *Alcanivorax*, *Alteromonas* and *Thalassiospira*, and phenanthrene degraders to the genera *Neptunomonas*, *Cycloclasticus* and *Colwellia*, whereas we found *Cycloclasticus* and *Colwellia* to be primary naphthalene degraders.

Both *Colwellia* and *Cycloclasticus* were non-detectable before enrichment, and were still rare after 24 hours of incubation with naphthalene (July, fig .4). However, after 88 hours of incubation they were the most dominant taxa in the mesocosms, and exhibited very significant enrichment indicating incorporation of naphthalene-derived carbon into their DNA over a substantial number of replication cycles.

The *Colwellia* MAG was the only one assembled here that contained an annotated subunit of naphthalene 1,2-dioxygenase (nahAb). As this is the first step of the pathway, requiring investment of reducing power [68], it is likely that at least some of the downstream process is also present in the complete genome. *Colwellia* was not abundant enough to detect a density shift after 24 hours, indicating that it may require a lag period in order to take full advantage of the naphthalene input, probably due to the initial low abundance of both *Colwellia* and *Cycloclasticus*. This could be explained if *Colwellia* uses naphthalene (and/or related compounds) as a sole carbon source as opposed to generalist taxa which use it in addition to other sources. This relates to *Colwellia* being normally rare (and likely sustained by consistent yet low inputs from atmospheric deposition and crude oil and diesel leaks), while the generalists such as *Planktomarina*, *Puniceispirillum* and SAR92 can be found in moderate abundances year-round.

*Cycloclasticus* genes dominated the metagenome of the phenanthrene-enriched DWH mesocosm [18], highlighting its importance as an obligate PAH degrader [69]. Unlike the *Colwellia* MAG, our *Cycloclasticus* MAG did not contain any part of the KEGG-defined aerobic naphthalene-specific degradation pathway which is based on pure cultures. However, it did include several ring-hydroxylating dioxygenases. While these enzymes are best known to degrade single aromatic hydrocarbons, previous studies demonstrated that some single-ring aromatic degrading enzymes are, in fact, capable of degrading naphthalene (two aromatic rings) efficiently [69–72]. Additionally, the *Cycloclasticus* MAG has a sigma-54-dependent transcription regulator with a potential hydrocarbon-binding domain as identified in *Cycloclasticus zancles* 78-ME (GenBank accession AGS40441.1), which could control transcription of hydrocarbon degradation genes. Dioxygenases require iron and an iron-binding domain, such as ferredoxin which can be shared by multiple enzymes [73, 74], which we found in this MAG. As we know that our *Cycloclasticus* strain can degrade naphthalene, and has genes that can participate in similar pathways but none of the traditional pathway, we posit that non-traditional degradation pathways were utilized to degrade naphthalene. It is reasonable to expect that generalist aromatics-degrading organisms might “most-economically” possess multi-functional pathways that do not fully optimize degradation of a single substrate but work adequately with several aromatic substrates, without the need to carry the “extra baggage” needed to optimize each specific pathway. To further support this idea, metatranscriptomes of the microbial community within the oil plume of the Deepwater Horizon spill revealed low to nonexistent transcription of known PAH-degradation genes despite their presence in metagenomes [75]. Moreover, the use of stable isotope probing indicated that both *Colwellia* and *Cycloclasticus*, and quite possibly the other six MAGs that were proven to be enriched, have the ability to degrade PAHs, whereas if only metagenomics was used we may have concluded that they did not.

Both *Colwellia* and *Cycloclasticus* displayed a potential for using a variety of nitrogen sources to different ends, with a full dissimilatory nitrite reduction to ammonium (DNRA) pathway and nitrite/nitrate transporters (FocA, NarK) as well as a urea transporter and the ammonium-assimilating glutamate synthase pathway. Nitrate, nitrite and ammonium are always detectable at POLA in surface seawater (sup. fig. S6), and thus are not limiting nutrients in this site. DNRA is an anaerobic process, which appears to be in conflict with the aerobic nature of surface seawater. However, PAHs are hydrophobic and tend to attach to particles, and particles can provide anaerobic microniches in their interior [76]. It is possible that PAH degradation occurs in large part on particles at POLA, which would explain the presence of this pathway within the MAGs, and that in these taxa nitrite is sometimes used as an oxidizer whereas ammonia and/or urea are used as nitrogen sources. PAH degradation on particles was also described above as a possible explanation for the labeling of picoeukaryotic autotrophs (some of which may also associate with the same particles).

*Litoricola* and *Porticoccus*, whose OTUs were enriched in this study, were previously shown to increase in relative abundance in oil-enriched mesocosm experiments [77]. While their genomes, as well as genomes of the other MAGs with matching enriched OTUs, reveal no full PAH or BTEX (benzene, toluene, ethylbenzene, xylene) degradation pathways, they do contain either aromatic ring hydroxylating genes or biphenyl dioxygenase. Biphenyl has a very similar chemical structure to naphthalene and biphenyl dioxygenase is known to degrade a variety of substrates [78], therefore it is possible that a variant of this enzyme can be used to degrade naphthalene.

While metagenomic read-recruitment and genes within MAGs only indicate a potential capability to degrade PAHs, a study in chronically contaminated Mediterranean sediments encompassing a wide selection of aromatic hydrocarbons revealed a positive correlation between abundance of genes coding for hydrocarbon degradation enzymes, 16S-rRNA of potential degraders and gas chromatography mass-spectrometry (GC-MS) of degradation rates and intermediates of degradation pathways [15]. This implies that in the case of PAH degradation processes presence of a pathway can be a fairly good indication of its expression given a substrate input.

Based on the functional pathways and ABC transporters found in MAGs of naphthalene incorporators, we propose targets for future experiments on enhancement of PAH bioremediation. As seven out of eight incorporator MAGs have a full glutamate synthase pathway, whereas only three have a full urea transporter, addition of ammonia rather than urea may be more beneficial. Similarly, phosphate is more likely to augment bioremediation than phosphonate. To supply iron for dioxygenase synthesis, adding heme/hemoproteins should be superior to addition of Fe(II) or Fe(III). Marine bacterioplankton have been shown to be able to incorporate iron from heme groups [79, 80].

In the current climate of excessive use of fossil fuels, chronic deposition of toxic and recalcitrant polycyclic aromatic hydrocarbons into the coastal ocean is inevitable. PAH-degrading bacteria may provide some control over the remineralization of these inputs and could serve as targets for bioremediation technologies. Identification of naturally-occuring biodegraders is a crucial first step, but optimization of the degradation process requires knowledge of the metabolic requirements of these organisms [21]. The combination of stable isotope probing with metagenomics could provide a source for genomically-generated hypotheses and experimental design for future bioremediation experiments.

## Supporting information

Supplementary Material Titles

## Acknowledgements

We extend our gratitude to Prof. Douglas Capone (USC) and Troy Eric Gunderson for the uptake rate measurements and for comments on this manuscript. We would like to thank Dr. Ben Tully for assistance with analysis of functional pathways. This research was made possible with the support of NSF grants 1136818 and 1737409, Gordon and Betty Moore Foundation Marine Microbiology Initiative grant GBMF3779 and The Wrigley Institute for Environmental Studies Norma and Jerol Sonosky fellowships to E.T.S.

