## Supplementary figures and images for "Metagenomics and stable isotope probing offer insights into metabolism of polycyclic aromatic hydrocarbons degraders in chronically polluted seawater"

### Supplemental Figure 1

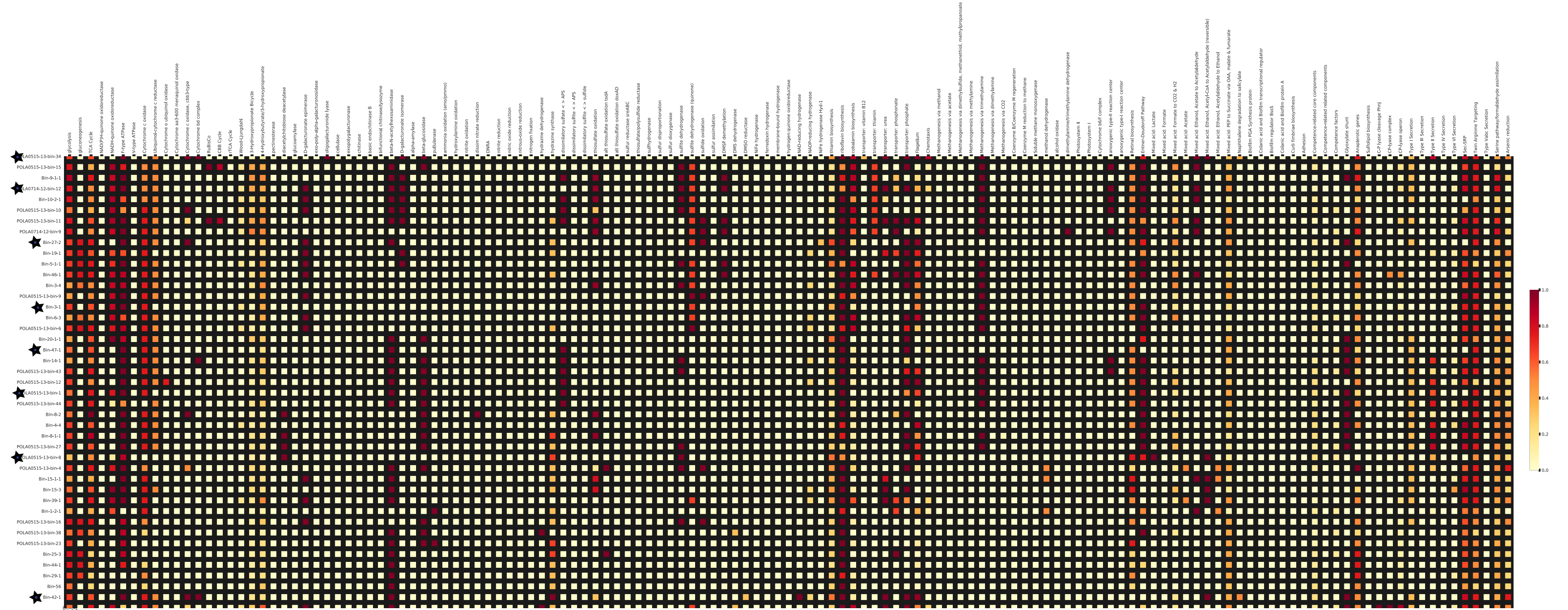

### Supplemental Figure 2

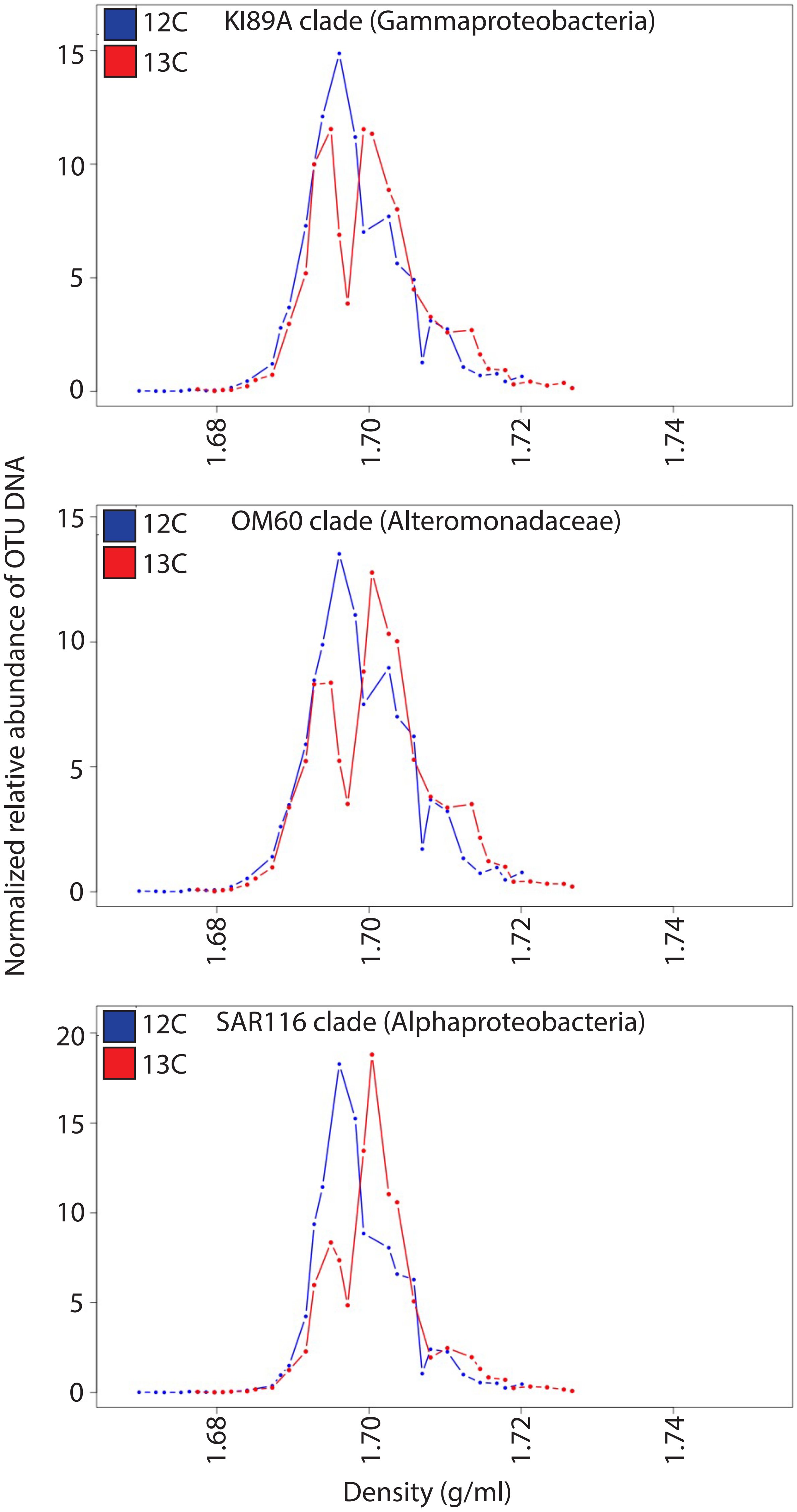

### Supplemental Figure 3

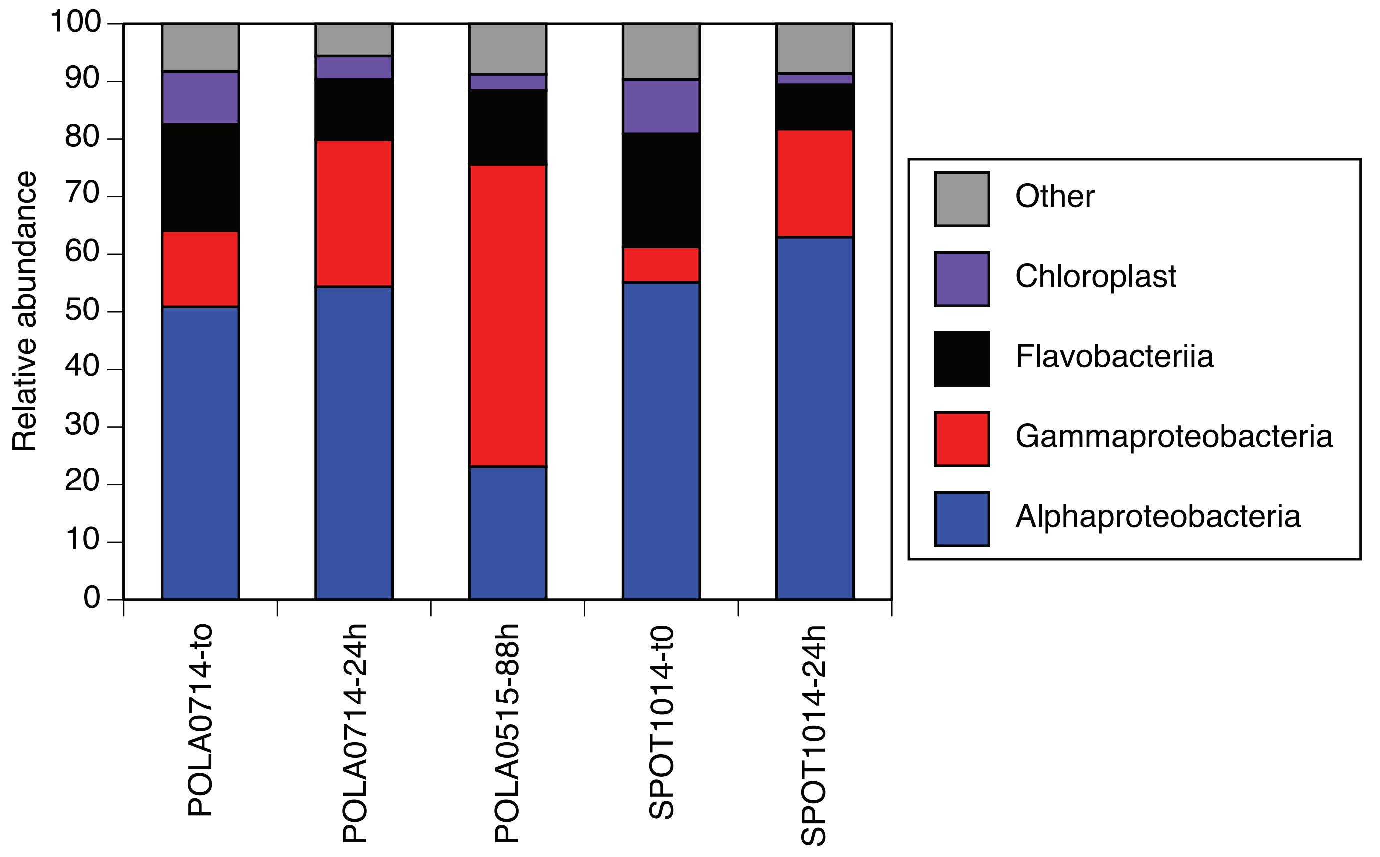

### Supplemental Figure 4

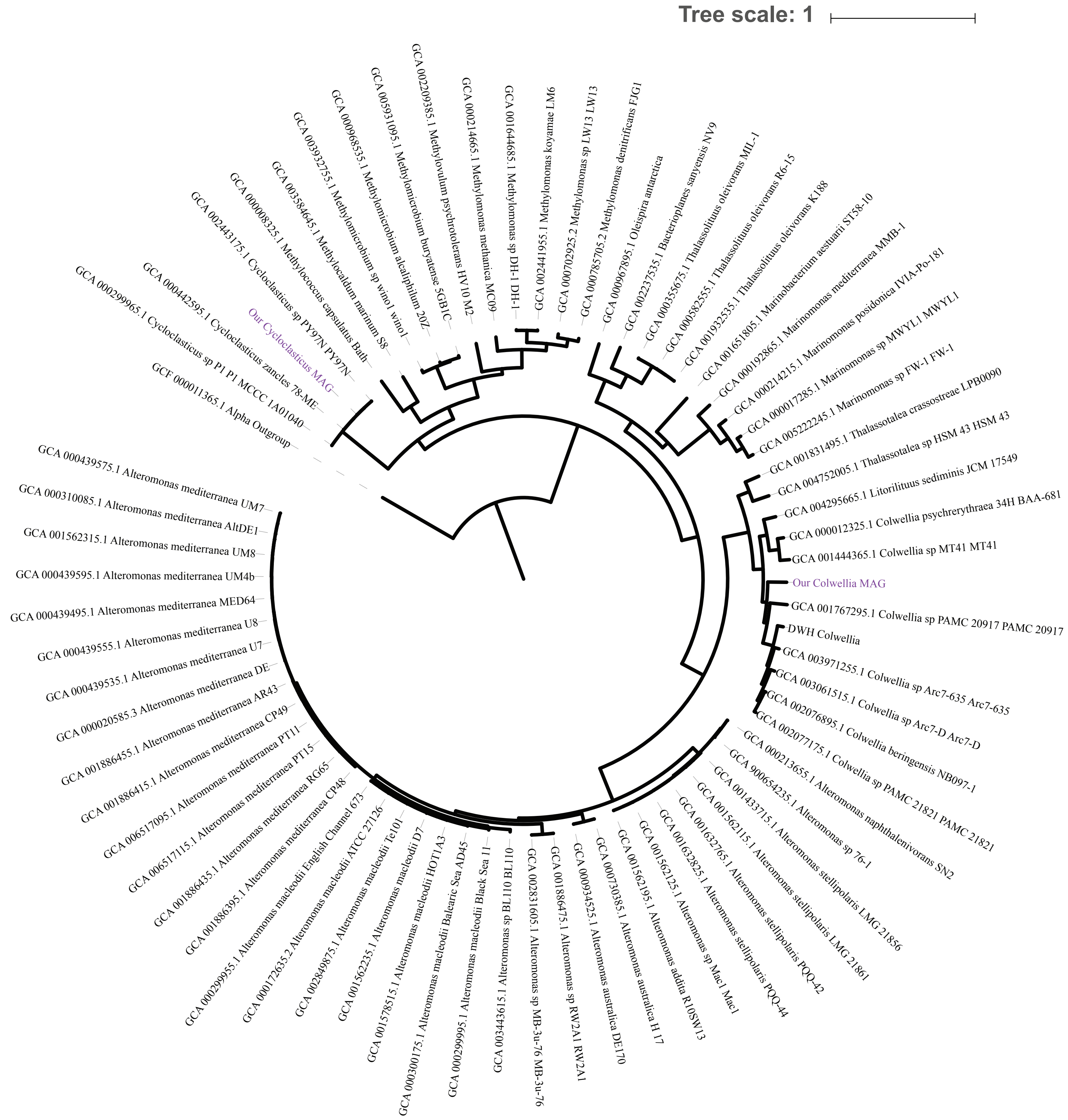

### Supplemental Figure 5

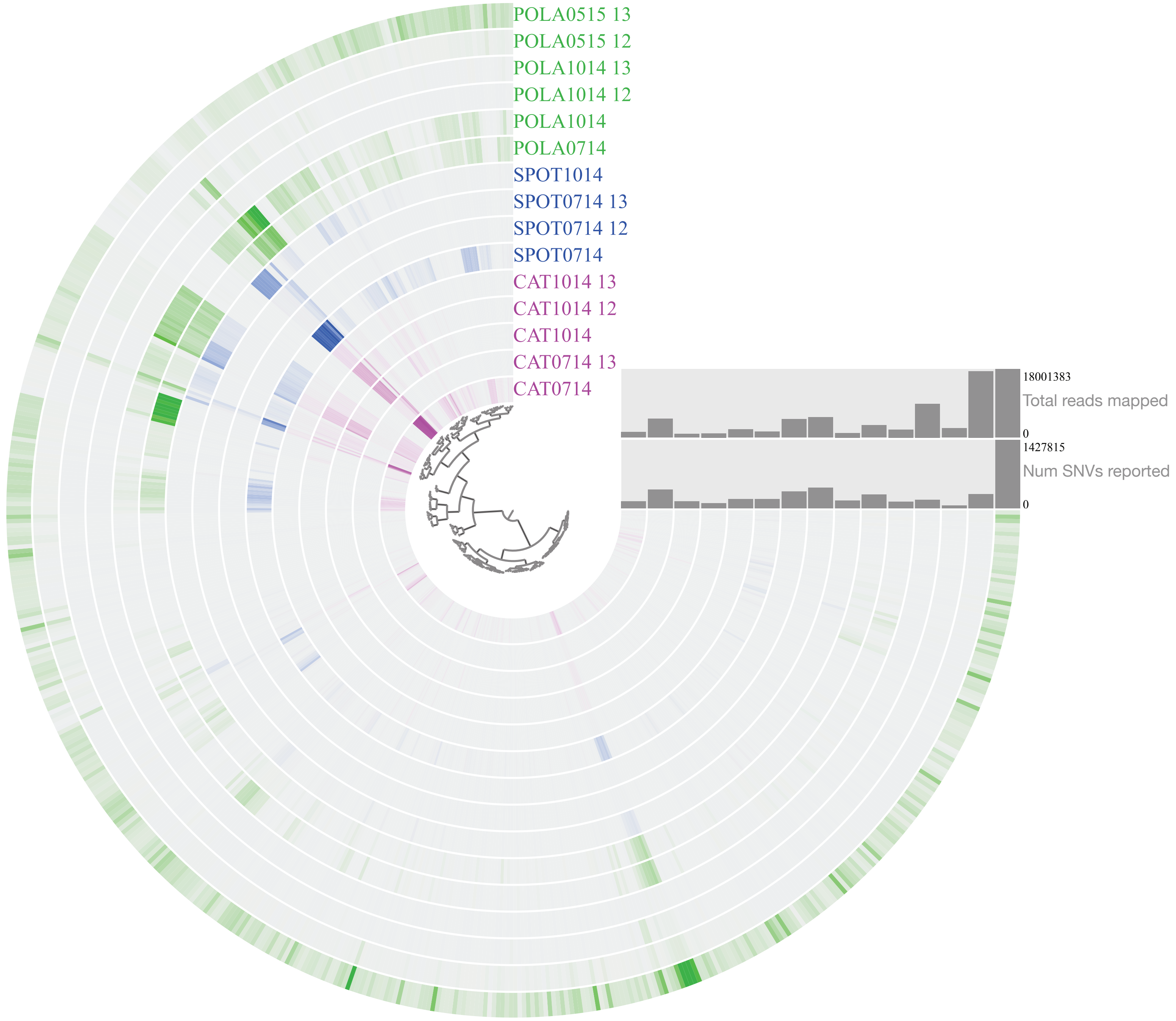

### Supplemental Figure 6

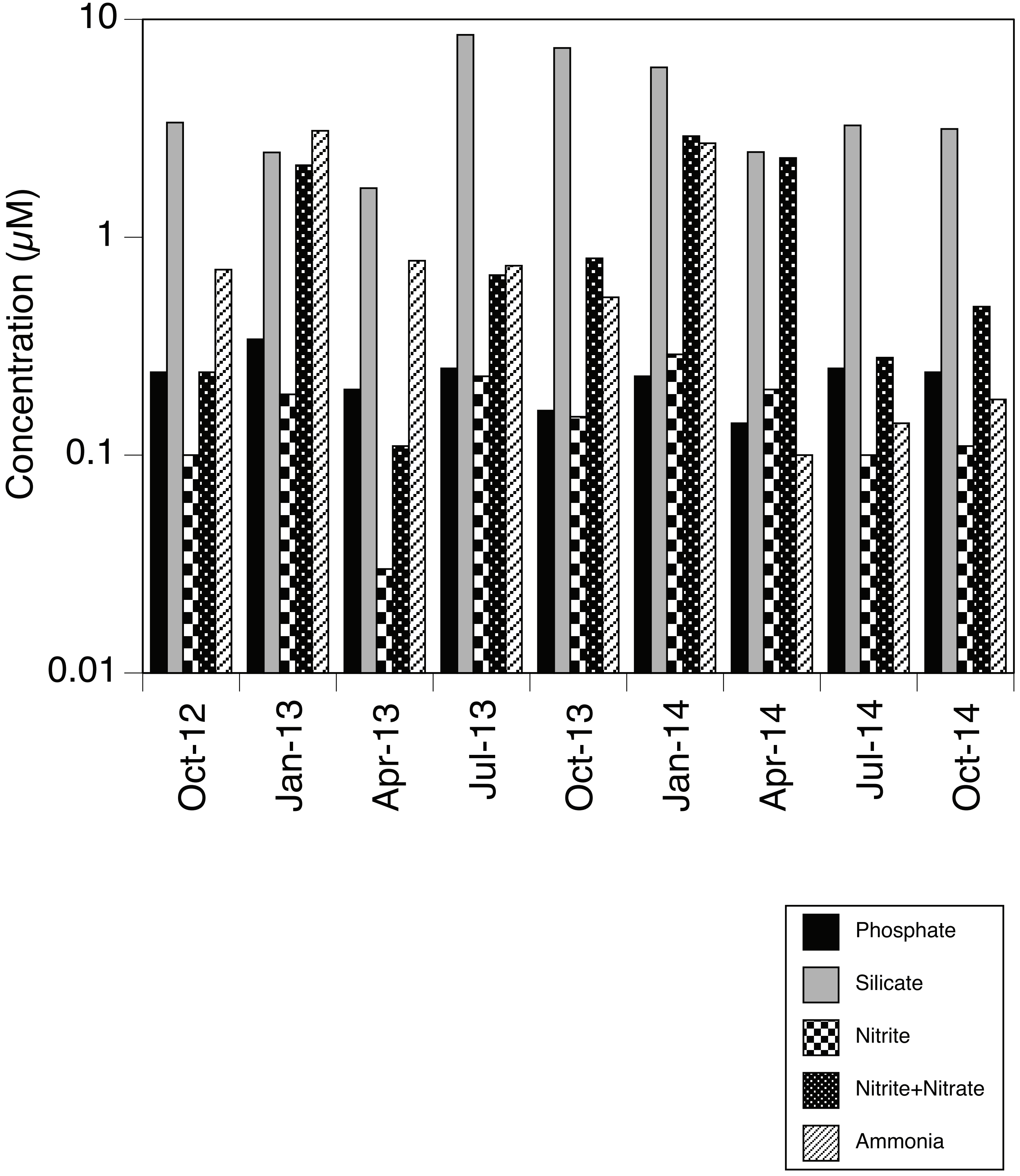
