## Supplementary Table 1 for "Metagenomics and stable isotope probing offer insights into metabolism of polycyclic aromatic hydrocarbons degraders in chronically polluted seawater"

| Sample | # MG raw reads | # MG post-QC paired reads | # post QC 16S-rRNA amplicons | Ultracentrifugation |
| --- | --- | --- | --- | --- |
| POLA 7/2014 12C | 51,205,975 | 44,358,529 | 476,467 | Yes |
| POLA 7/2014 13C | 58,167,816 | 52,822,155 | 1,450,317 | Yes |
| POLA 7/2014 t_0_ | 26,315,915 | 22,764,173 | 15,741 | No |
| POLA 10/2014 12C | 20,875,980 | 14,067,988 | N/A | No |
| POLA 10/2014 13C | 14,297,951 | 8,159,916 | N/A | No |
| POLA 10/2014 t_0_ | 26,952,250 | 23,355,914 | 26,570 | No |
| POLA 5/2015 12C | 27,623,601 | 24,760480 | 841,462 | Yes |
| POLA 5/2015 13C | 49,018,041 | 32,827,498 | 1,710,651 | Yes |
| SPOT 7/2014 12C | 24,465,312 | 21,436,796 | N/A | No |
| SPOT 7/2014 13C | 30,268,423 | 26,493,414 | N/A | No |
| SPOT 7/2014 t_0_ | 26,267,706 | 17,918,445 | 15,532 | No |
| SPOT 10/2014 12C | No metagenome | No metagenome | 468,340 | Yes |
| SPOT 10/2014 13C | No metagenome | No metagenome | 339,265 | Yes |
| SPOT 10/2014 t_0_ | 29,795,731 | 20,495,706 | 24,905 | No |
| CAT 7/2014 13C | 27,318,562 | 23,216,992 | N/A | No |
| CAT 7/2014 t_0_ | 25,234,021 | 17,223,728 | 40,936 | No |
| CAT 10/2014 12C | 24,098,874 | 21,481,089 | N/A | No |
| CAT 10/2014 13C | 26,675,357 | 22,882,709 | N/A | No |
| CAT 10/2014 t_0_ | 25,234,021 | 18,632,041 | 15,724 | No |
