## Supplementary Material Titles for "Metagenomics and stable isotope probing offer insights into metabolism of polycyclic aromatic hydrocarbons degraders in chronically polluted seawater"

Supplementary figures titles:

S1: Basic metabolic pathways in MAGs represented as % of the enzymes present. MAGs matching enriched OTUs are labeled with a star.
S2: Bimodal density shifts in OTUs from SPOT sample from October.
S3: Changes in relative abundance of taxonomic groups between t0, 24 hours and 88 hours
S4: Phylogenomic tree showing the placement of our *Colwellia* and *Cycloclasticus* MAGs.The MAGs are labeled in purple. The tree is based on 117 single-copy genes of Gammaproteobacteria and was built in GToTree. Complete reference genomes were downloaded from RefSeq on 8/20/2019.
S5: Mean coverage (Q2Q3) of viral contigs in naphthalene-enriched and t0 metagenomes. Each concentric circle represents a sample. Each bar stands for one viral contigs. The color intensity correlates to mean coverage. The central tree represents hierarchical clustering of the viral contigs by tetranucleotide frequency and mean coverage.
S6: Nutrient concentrations at the Port of Los Angeles

Supplementary table titles:
S1: Sample processing details – number of metagenomics (MG) reads, 16S-rRNA amplicons and ultracentrifugation. Number of amplicons in enriched mesocosms (not t0) are combined across all fractions.
S2: KEGG annotations and best blast hits (NCBI NR) to open reading frames identified in the metagenomic assembled genomes.
S3: General information on metagenomic assembled genomes (% completion, % redundancy, length) and taxonomy determined by GTDB-Tk, Anvio, GToTree and 16S-rRNA
S4: KEGG pathways and transporters identified in the metagenomic assembled genomes
